# Understanding How Biopolymers Found in Marine Ecosystems Impact the Efficacy of Bacteriophages against Bacteria

**DOI:** 10.1101/2024.11.07.622396

**Authors:** Rhea Rupareliya, Joyce Kwack, Dylan Quan, Tay Kim

**Affiliations:** San Diego State University

## Abstract

Bacteriophages are abundant in Earth’s ecosystem and serve as valuable mediators in marine environments, regulating bacterial populations in oceans, hot springs, and salterns, among others. However, bacteriophages have high-specificity to their host, thus making it difficult to study bacteria-phage relationships. In this study, we observe whether naturally occurring biopolymers (coral mucin (from Stylophora pistillata), chitin, and sodium alginate) increase bacteriophage affinity to bacteria by recording the plaques formed throughout 15-minute time intervals. We use an isolated *E. coli* strain and its corresponding T7 phage, as well as isolated bacteria and phage strains from a coral sample (labeled Stylophora pistillata or “Coral 1”). The research conducted plaque assays using minimal media solution to quantify phage-forming units (PFUs) using 2 bacterial forms, Coral 1 and E. coli and Mucin, Sodium Alginate, and Chitin as biopolymers. Our research found that the biopolymers had an effective impact on the generation of PFUs in our bacterial cultures for all samples. While we were unable to conduct future plaque assays, the information we have now is important to discovering ways to increase phage efficiency in marine environments to help maintain nutrient resource levels in such ecosystems.

## Introduction

Bacteriophages are essential to microbiomes in order for bacterial regulation and are key to increasing nutrient availability in marine ecosystems [1]. These viruses are also critical to increase host gene variation through horizontal gene transfer, making them critical to increasing diversity in marine environments [2]. Even though bacteriophage are integral to marine environments, there still exists a lack of in-depth research on bacteria-phage relationships.

While bacteriophage have superior prolific tendencies in comparison to bacteria, as phage produce 100-200 new phage particles during reproductive cycles while bacteria only produce 2 daughter cells,phage are still unable to outnumber bacterial host populations [3]. Yet, 20-40% of bacteria are eliminated by these phages in marine ecosystems–thus, we believe it is important to increase knowledge of bacteria-phage relationships to help modulate bacteria in marine ecosystems. In fact, bacteriophage are important in harmonizing the bacteria population by increasing abundance and diversity, as well as changing physiology, competitive ability, and virulence [3].

Phage are crucial to the turnover of organic material, as phage-mediated lysis releases organic matter into the ecosystem and therefore creates an essential nutrient supply for both the host and the environment [9, 10]. They also ensure that bacterial populations do not overpopulate and dominate in an ecosystem, since they are great at supporting diversity and preventing too much overflow of certain bacterial species. [11, 12]. This diversity is crucial to supporting the health of the ecosystem and ensuring proper growth in marine environments [13,14].

Unfortunately, there is very limited details and research on phage functions in the coral and marine ecosystem since it is difficult to formulate viral genomes and bacterial hosts from coral samples [15]. At the reef scale, phages are vital to creating competitive dynamics in the benthic space and biogeochemical cycling which are important for reef growth [16, 17]

In the past, *Vibrio mediterrane*, a pathogen that infects coral species has been observed when studying bacteriophage-bacteria interactions, and research has found that two phage species exist in the coral mucus layer, providing an added layer of defense against this pathogen [18,19]. Phage have shown in the past to be very valuable treatments for bacterial infections in coral environments.

As antibiotic resistance becomes an increasingly prevalent threat to the viability of microbiomes, it is vital that we find a new solution. In that regard, bacteriophage presents a capable and promising strategy to minimize antibiotic resistance, while also managing bacterial populations. [20].

## Methods

### 1. Bacteria isolation

1. In order to make 100 mL of top agar solution, add 2.5 g of tryptone, 0.5 g of yeast, 0.5 g of NaCl, and 0.65 g of agar to a sterile glass bottle.
2. In order to make 500 mL of the agar plate solution, add 5 g of tryptone, 2.5 g of yeast, 2.5 g of NaCl, and 7.5 g of agar to a sterile glass bottle.
3. Add distilled water to both bottles until it reaches the desired volumes.
4. Autoclave at 120° C.
5. Obtain sample and add 1 mL of sample and 9 mL of phosphate buffered solution to a container.
6. Repeat with 1 mL of the resulting mixture and 9 additional mL of phosphate buffered solution for a second serial dilution.
7. Obtain up to 10 Petri dishes and a Bunsen burner.
8. Label the plates.
9. Bring the agar solutions back from the autoclave. Cool with water if necessary.
10. Pour a thin layer of the agar plate solution into each one, using a sterilized metal loop to get rid of any bubbles.
11. Allow the agar to dry.
12. Add a thin layer of the top agar solution to at least 3 plates. If the top agar solution has hardened, microwave until liquid again.
13. Streak the sample solution onto 2 plates once the top agar is dry using the quadrant method. Have 1 control plate streaked only with a buffer. Sterilize the rim of the containers and the metal loop with the flame.
14. Clearly label the Petri dishes, invert them, and incubate overnight.
15. After incubating the plates, check for bacterial growth.
16. Grab a colony and check growth to make sure it is the same bacteria.
17. Use pipette tip to grab a single colony and inoculate them in a buffer solution.
18. Label isolated bacteria (“C1” in the case of our experiment).

## 2. Phage Isolation

### 2.1 PEG Precipitation of Viral Concentrates

2.1.1 Collect 1mL of sample for microscopy if needed. Follow microscopy protocol for VMR quantification.
2.1.2 Add 10% PEG 8000, and 5% NaCl to a conical tube(s). Add water sample and dissolve. Be sure to modify the amount of salt added for marine samples. Check refractometer reading for calculation. Final NaCl should be 5%. Modify the amount of PEG 8000/NaCl added as needed per sample volume. Most samples will be 35-40 ppm but salterns might have more… For samples >250mL: use 250mL conical tubes, duplicates may be prepared depending on sample volume. For samples <100mL: use 50mL conical tubes.
2.1.3 Clearly label samples. Let samples settle overnight at 4°C.
2.1.4 Centrifuge at 3,000rpm for at least 14 hrs. Alternately, you may spin at 11,000xg at 4°C for 40 min using the RC5C centrifuge (rotor code 3, brake off).
2.1.5 If needed, save 2mL of supernatant per sample.
2.1.6 Discard remaining supernatant making sure the invisible pellet is not disturbed.
2.1.7 Place inverted tubes on Kimwipes to remove droplets, let rest.
2.1.8 Resuspend pellet with 5mL SM buffer, vortex. For samples with multiple tubes add enough buffer for a combined resuspension volume of 5mL. Ex. A sample of 500mL split into 2, 250mL conical tubes, resuspend with 2.5mL of SM buffer in each tube.
2.1.9 Transfer resuspended pellet into a labeled 5mL tube and proceed to CsCl isolation protocol.

### 2.2. Cesium Chloride Sample Preparation of Phage Community DNA

2.2.1 Make phage concentrates (lysate or filtered water sample)
2.2.2 Set up CsCl gradient shown in figure below. Make gradients from the same buffer samples are in. Make sure the solution used to make gradients have been filtered through a 0.02um filter. Mark outside of the tube to denote the location of each fraction. Note: Maximize the number of tubes that can go into the SW41 Ti rotor - 6. **BALANCE EXACTLY BEFORE CENTRIFUGING** (±0.001g)
2.2.3 Centrifuge at 22,000rpm at 4°C for 2 hrs. using the SW41 Ti. Make sure to write the ending counter number in the logbook. (ALT. RM 409, Beckman IE-80, 30000rpm)
2.2.4 Pierce the tube with a 16 gauge needle at the red arrow and pull ∼1.5mL into a syringe, collecting the fraction and boundary between 1.35g/mL and 1.5g/mL
2.2.5 Transfer phage into a sterilized tube and store

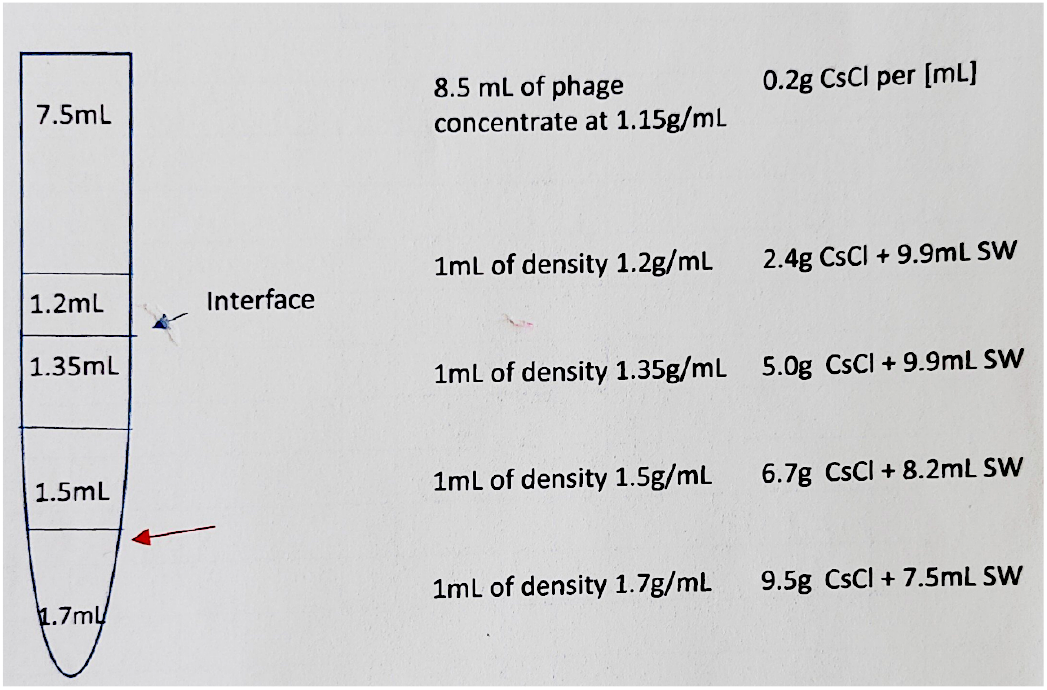

## 3. Prepare the Sugar Solution (Coral Mucin)

1. Ensure the corals are in good health before starting the experiment.
2. Dissolve the desired sugar (e.g., sucrose or glucose) in filtered seawater to create a stock solution. Common concentrations range from 5 mM to 50 mM.
3. Sterilize the sugar solution by filter sterilization using a 0.22 μm filter.
4. Place coral fragments in individual containers filled with filtered seawater.
5. Add the sugar solution to the containers to achieve the desired final concentration. For example, if using a 10 mM final concentration, calculate the volume needed based on the container’s water volume.
6. Ensure thorough mixing of the sugar solution with the seawater.
7. Incubate the corals in the sugar-treated seawater for a specified period, usually between 24 to 48 hours.
8. Maintain the corals in a controlled environment with stable temperature, light, and aeration.
9. After the incubation period, observe the corals for increased mucus production.
10. Collect the mucus using a sterile pipette or by gently shaking the coral fragments in the water and then pipetting the mucus-rich water.
11. Transfer the coral fragments back to their original tank with clean filtered seawater.
12. Monitor the corals for recovery and health status.

### Mucus isolation protocol

1. Empty out mucus-filled-water produced by coral fragments. Filter out new water using a .22 um from the aquarium tank to replace and change.
2. Fill mucus/water solution in 50 mL or 250 mL Eppendorf tubes. Centrifuge at atleast 4,000 rpm for 15 minutes to create a mucus pellet filled with sediment.
3. Pour out supernatant into a beaker or graduated cylinder. Discard the tube containing the sediment pellet.
4. Gradually pour in 2x volume of isopropyl alcohol (2-propanol) into the beaker. Mix gently and continue to add more until 2x volume of isopropyl alcohol is reached. (There should begin to be a cloudy color change).
5. Wait 30 minutes to several hours to allow the mucin glycoproteins to precipitate with isopropyl alcohol.
6. Gently pour the mucin-alcohol solution back into Eppendorf tubes. Centrifuge at around 1,500 rpm for 4 minutes.
7. Discard the supernatant and fill with a buffer (PBS, etc.).
8. Transfer solution into a maximum surface area container.
9. Place under the UV fume hood for at least 1 hour.
10. Extract and place mucin back into a tube. Fill with PBS buffer
11. Wash 2x with PBS buffer to remove the alcohol dissolved within the mucin
12. Mix with minimal media and proceed to plaque assay.

### 4.1 L Minimal Media Preparation

1. To prepare 1 liter of M9 minimal media, the following salts are required:
  a. Na2HPO4 (Sodium phosphate dibasic) – 33.9 g
  b. KH2PO4 (Potassium phosphate monobasic) – 15.0 g
  c. NaCl (Sodium chloride) – 2.5 g
  d. NH4Cl (Ammonium chloride) – 5.0 g
2. Dissolve the Na2HPO4, KH2PO4, NaCl, and NH4Cl in approximately 900 ml of distilled water. Adjust the final volume to 1 liter.
3. Autoclave the M9 salt solution in 120°C for approximately 30 to 40 minutes
4. After the solution has cooled, aseptically add the following sterile components:
  a. 1 M MgSO_4_ (Magnesium sulfate): 1 mL
  b. 1 M CaCl_2_ (Calcium chloride): 0.1 mL
  c. Carbon source (e.g., glucose, glycerol): 20 g

## 5. Plaque Assay Preparation

1. For each trial, measure the appropriate quantities in sterile 15 mL corning Falcon tubes
  a. Control: 10 mL of minimal media
  b. Chitin: 10 mL of minimal media + 0.01 grams of chitin powder
  c. Sodium Alginate: 10 mL of minimal media + 0.005 grams of sodium alginate powder
  d. Mucin: 9 mL of minimal media + 1 mL of 5% mucin solution
2. Label the tubes accordingly
3. For each trial, add in the polymers
  a. Control - nothing
  b. Chitin - 0.01 grams of chitin powder
  c. Sodium Alginate - 0.005 grams of sodium alginate powder
  d. Mucin - 1 mL of 5% mucin solution
4. Bacterial and Phage Inoculation Each trial is performed using a specific bacterial strain and its corresponding phage. The bacteria must be inoculated in Luria Broth for about 2 hours before use in order to ensure it is in the proper phase. The bacterial strains and phage pairings are as follows:
  a. Trial 1 (E. coli):
    i. Bacterial strain: E. coli (mid-log phase)
    ii. Phage: T7 phage
    iii. Conditions: Control, Chitin, Sodium Alginate, Mucin
  b. Trial 2 (C1 bacteria):
    i. Bacterial strain: C1 (mid-log phase, isolated from previous experiment)
    ii. Phage: Phage isolated from C1 bacterium
    iii. Conditions: Control, Chitin, Sodium Alginate, Mucin
  c. Trial 3 (Saltern bacteria):
    i. Bacterial strain: Saltern (mid-log phase, isolated from marine environment)
    ii. Phage: Phage isolated from Saltern bacterium
    iii. Conditions: Control, Chitin, Sodium Alginate, Mucin
  d. Trial 4 (C2 bacteria):
    i. Bacterial strain: C2 (mid-log phase, isolated from previous experiment)
    ii. Phage: Phage isolated from C2 bacterium
    iii. Conditions: Control, Chitin, Sodium Alginate, Mucin
5. For each trial, introduce the bacterial culture into the Falcon tubes containing the respective minimal media and polymer conditions. Maintain consistent bacterial concentrations across all samples. Inoculate each tube with a known volume of the corresponding phage stock, ensuring equal phage concentrations for reliable comparison between control and polymer-treated conditions.

## 6. Sampling and Plating

1. Every 15 minutes, take small samples (e.g., 1 ml) from tubes.
2. Immediately transfer each sample to a sterile centrifuge tube.
3. Centrifuge the samples at an appropriate speed (e.g., 10,000g for 5-10 minutes) to pellet bacteria and phage complexes.
4. After centrifuging, carefully discard the supernatant and keep the bacterial/phage pellet.
5. Resuspend the pellet in a small volume of sterile minimal media (e.g., 100 μl).
6. Mix 100 μl of resuspended pellet with 100 μl of fresh mid-log bacteria.
7. Add this mixture to 3 ml of molten top agar (cooled to ∼50°C) and immediately pour it over a base agar plate.
8. Let the agar solidify
9. Clearly label the Petri dishes and invert them

## 7. Incubation and Plaque Counting

1. Incubate the plates at the optimal temperature for the bacterial strain used (e.g., 37°C for E. coli).
2. Check for plaque formation after 4-8 hours or the following day, depending on the bacterial-phage system.
3. Continue sampling and plating at 15-minute intervals (e.g., 15, 30, 45, and 60 minutes).
4. Ensure that enough plates are prepared to cover all time points for both control and polymer-treated samples.
5. After incubation, count the number of plaques formed on each plate and record the time to successful plaque formation.

## 8. Data Analysis

1. Compare the number of plaques and the time to plaque formation between control and polymer-treated samples for each bacterial-phage system. Assess whether the presence of polymers (chitin, sodium alginate, or mucin) accelerates or hinders plaque formation compared to the control across the different bacterial strains and phage combinations.

## Results

### E. COLI

**Control (Fig 1.1):**
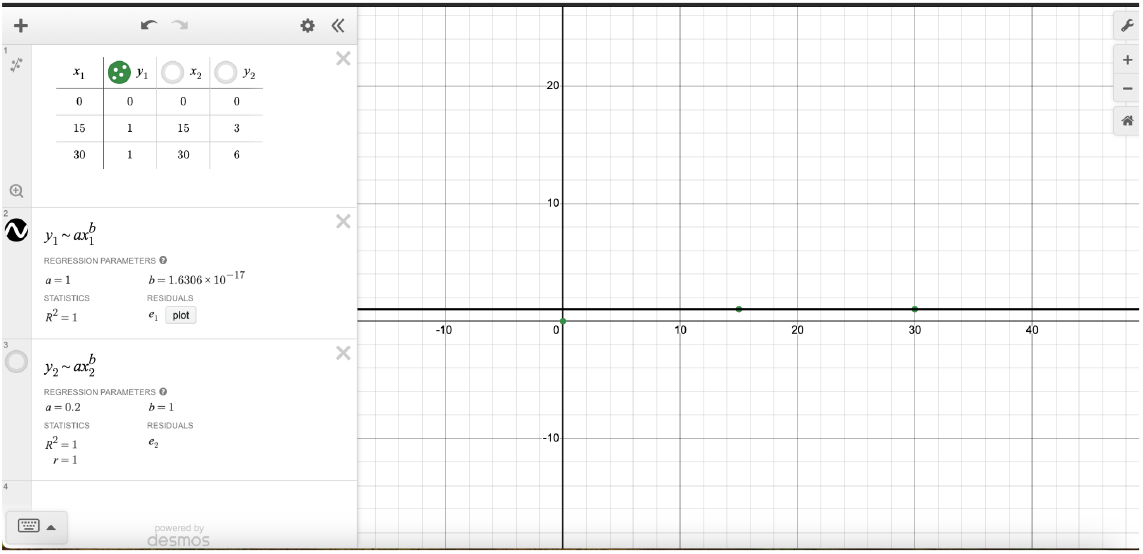
Graph of time versus PFUs for control group

**Chitin (Fig 1.2):**
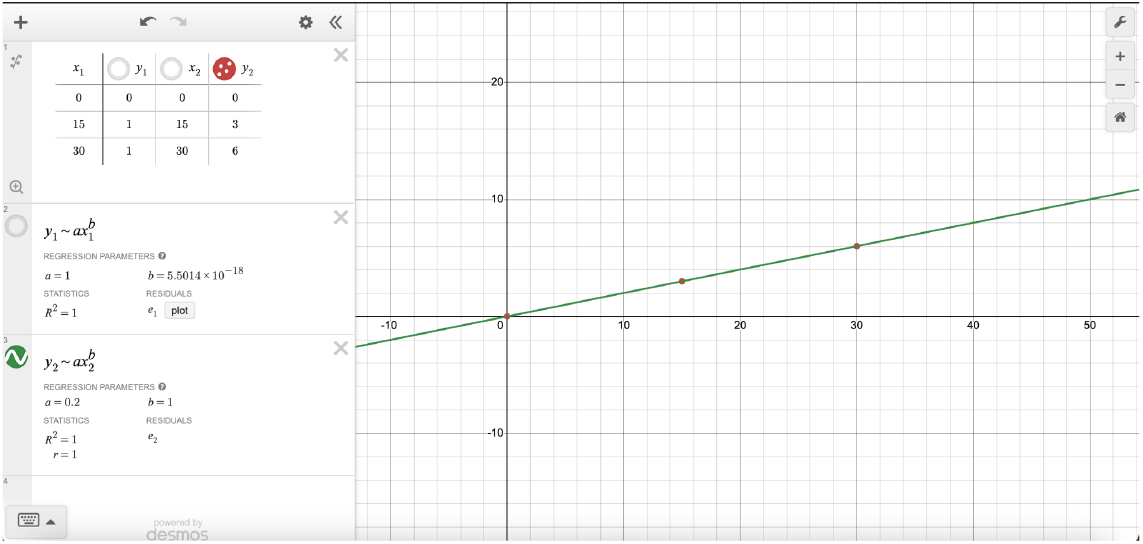
Graph of time versus PFUs for chitin group

### Stylophora pistillata

**Control (Fig 2.1):**
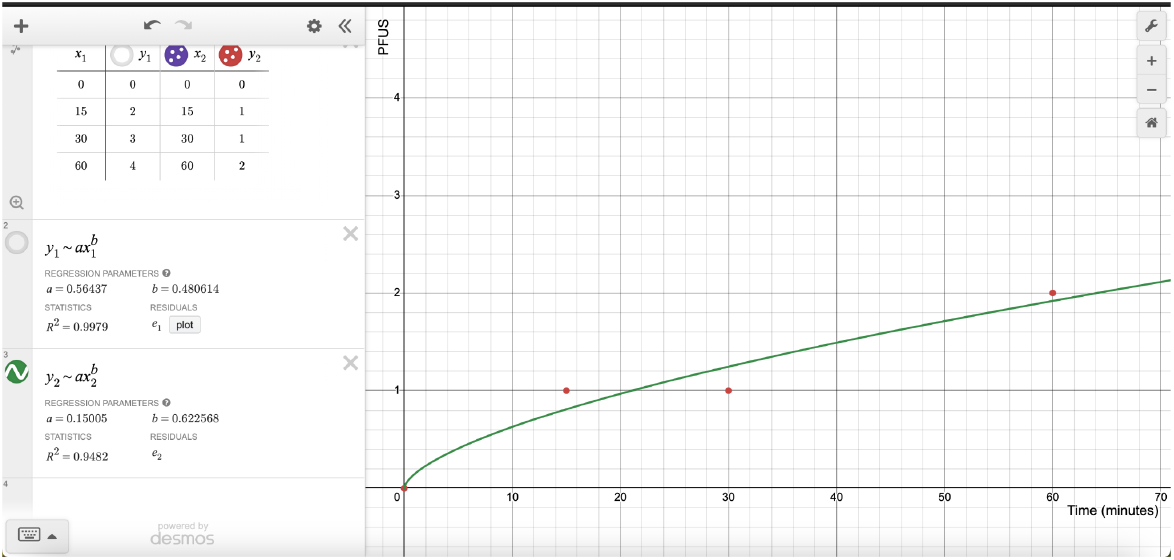
Graph of time versus PFUs for control group

**Chitin (Fig 2.2):**
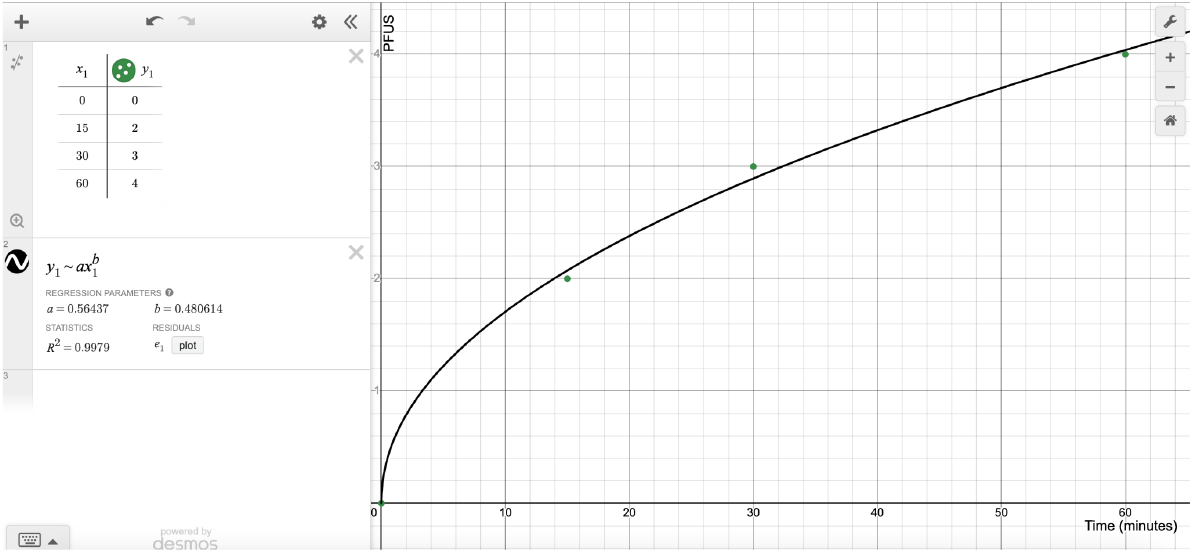
Graph of time versus PFUs for chitin group

**Coral Mucin (Fig 2.3):**
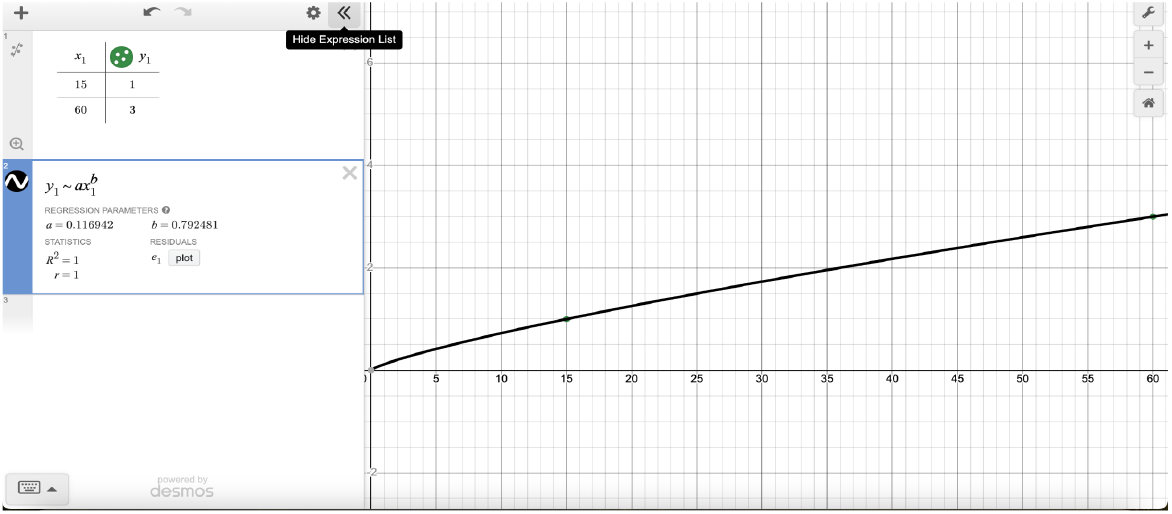
Graph of time versus PFUs for coral mucin group

**Sodium Alginate (Fig 2.4):**
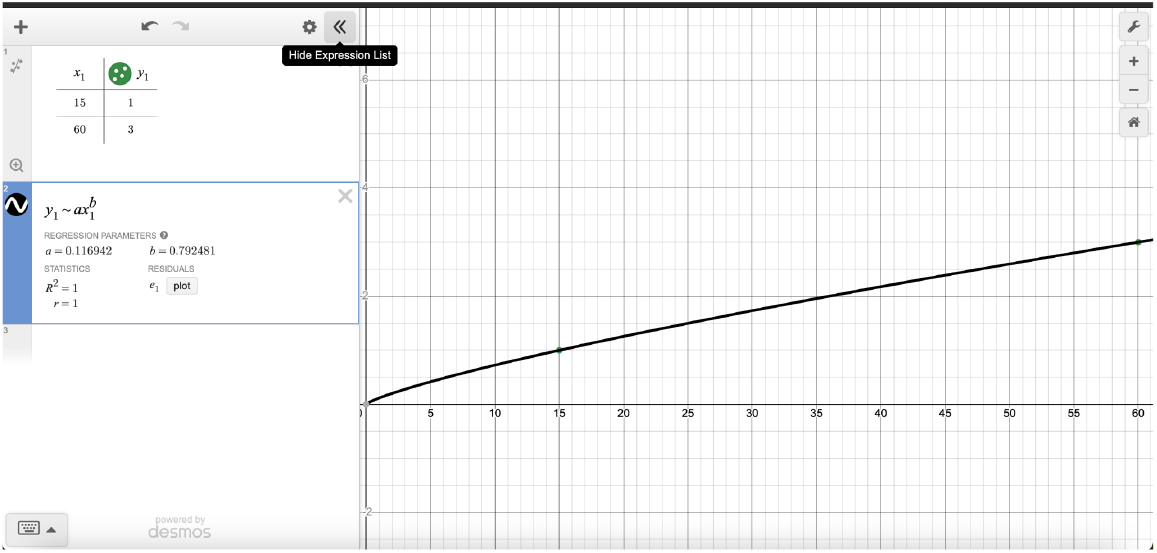
Graph of time versus PFUs for sodium alginate group

### Stylophora pistillata

**Fig (3.1):**
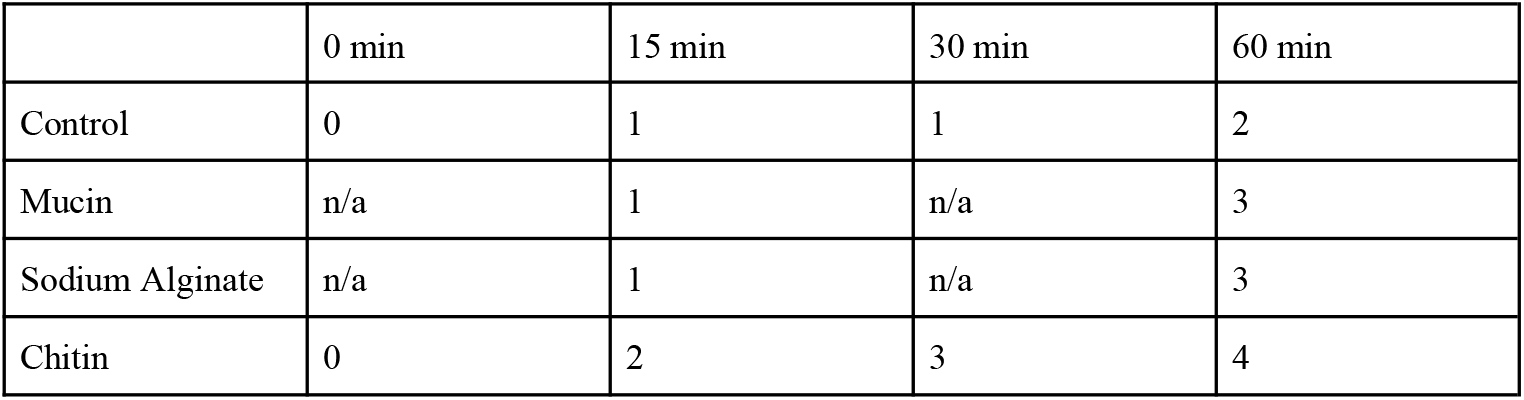
Table of values for different time intervals among the different groups for coral sample

### E. Coli

**(Fig 4.1):**
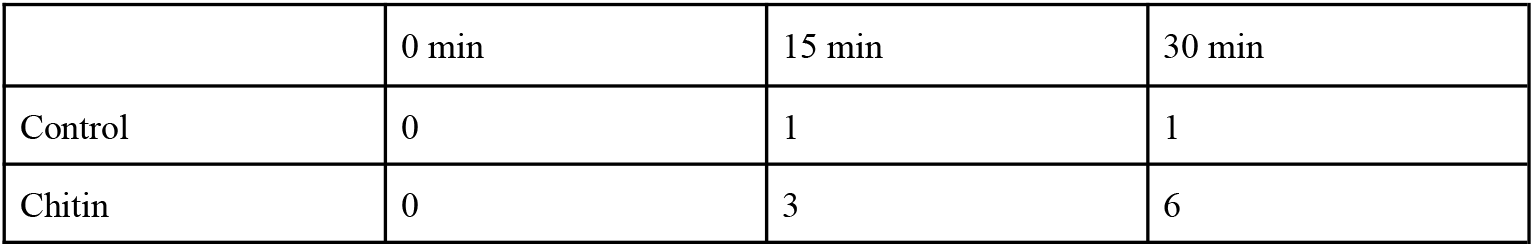
Table of values for different time intervals among the different groups for coral sample

We observed that polymers increased the rate at which PFUs, or plaque-forming units, appear within the agar plates. With a phage known to attack E. coli, T7 phage, we noticed that the presence of any polymers helped to facilitate virus-bacteria interactions due to an increase in the number of plaques formed that appeared as time went on. For this specific trial, the experiment was conducted for 30 minutes with both solutions kept in the incubator for 30 minutes. Only 1 plaque formed (Figure 1.1 and 3.1) on the agar plate without any polymer - the control group - after 30 minutes of incubation compared to chitin, the second most abundant biopolymer and a structural polysaccharide [21], which had around 6 plaques formed (Figure 1.2 and 3.1).

In another trial conducted using a bacteria strain produced by a Stylophora pistillata coral fragment and a suspected phage, a similar pattern emerged. After adding 8 g of C_11_H_22_O_11_, or sugar, to 240 mL of water containing the coral fragment, we allowed the coral fragment to produce an excess amount of mucus; the solution would ultimately be the bacteria used to conduct this set of trials and the coral fragment was placed into another container full of filtered sea water. The same procedure was executed. After 60 minutes, only 2 plaques formed on the agar plate containing no polymer, or the control set (Figure 2.1 and 4.1). Additionally, another biopolymer was used in this second trial: sodium alginate. Sodium alginate —a natural, water-soluble polysaccharide typically found in algae or seaweed that is known to be a thickening agent [22,23]— only furthered our conclusion as 3 PFUs also appeared on its agar plate after 60 minutes of incubation (Figure 2.2 and 4.1).

For the mucin plate, there were 3 plaques formed (Figure 2.3 and Figure 4.1). To extract the mucin, which was dissolved within the water, we mixed in isopropyl alcohol to precipitate mucin-clumps. The solution was then centrifuged and the mucin was collected after UV sterilization. The mucin was then used as another biopolymer after mixing the mucin into our minimal Media. For chitin, a total of 4 PFUs appeared on its plate (Figure 2.4 and 4.1).

Altogether, both trials and their concluding results indicate that biopolymers do increase the rate in which bacteriophages locate and bind to their host bacteria strain. Our studies demonstrate that biopolymers facilitate the attachment of bacteriophages known to attack a specific host (T7 phage). With phage that is suspected or hypothesized to attach to a specific bacteria colony, we also found that biopolymers improved the rate at which plaques formed. Thus, biopolymers can be used to essentially “attract” and allow for phages to attach to different bacteria strains. While bacteriophages are ubiquitous and abundant in nature, our hypothesized mechanism involving polymers can only further our knowledge of the unique nature of phage.

Trial 1:

Domain restriction of 0 to 30 minutes

E. coli

Chitin PFU rate: y=.2x

Control PFU rate: y=1

Trial 2:

Domain restriction of 0 to 60 minutes

Stylophora pistillata

Control: y=.15005x^.622568^

Chitin: y=.56437x^.4608^

Mucus: y=.116942x^.792481^

Sodium Alginate: y=.116942x^.792481^

## Discussion

In order to evaluate the validity of our hypothesis, we must outline the expected results. If polymers do bolster the ability of bacteriophages to find and lyse bacteria, then we would expect plaques to form significantly earlier or more frequently in the assays where the phage was included in the initial minimal media solutions, compared to the controls. If polymers have no effect on bacteria-phage relationships, then we would expect similar results regardless of polymer presence. In both bacteria strains, plaques started forming after the solutions had been incubating for 15 minutes. There was a noticeable difference in the number of plaques that formed with and without chitin for *E. coli*. This suggests that chitin did increase the T7 phage affinity for *E. coli*, though further trials may be necessary to confirm this. The difference was less noticeable for C1 bacteria, but the number of plaques formed overall for groups with the polymer was greater than the number of plaques formed in the control.

Based on our results, we can conclude that biopolymers, especially chitin, may improve the ability of phages to find and lyse bacteria. For the coral bacteria, sodium alginate and mucin appeared to have some effects on plaquing success and speed, but not as noticeable of an effect as chitin. This effect may be even more significant in natural marine environments, where chitin, sodium alginate, and coral mucin are naturally found. The addition of synthetic polymers may also greatly impact phage-bacteria interactions by also enhancing phage penetration and attachment to specific host bacteria [24].

Additional trials may corroborate and further this finding. It will possibly be beneficial to perform the protocols with various strains of the same bacteria, especially *E. coli* due to its accessibility, as well as with various types of bacteriophages that infect the same bacteria to see if results differ between tests. It would be interesting to also test different polymers, especially ones from other aquatic environments, such as extracellular polymeric substances (EPSs) [25]. Future experiments can test different pH levels, temperatures, and ionic strengths to assess if other factors can also affect phage-bacteria interactions; changing the concentration of phage to bacteria cells can also be evaluated for additional effects on phage-and-bacteria-interactions. More experiments and future trials will only aid our world’s understanding of the mechanisms that bacteriophages use to operate and lyse bacteria.

## Acknowledgements

We would like to thank Dr. Forest Rohwer, Andres Sanchez Quinto, Nelson Vayda and Vanessa Salcido for allowing us to use their lab space and serving as mentors for this project.

## References

[1] Brown, T. L., Charity, O. J., & Adriaenssens, E. M. (2022). Ecological and functional roles of bacteriophages in contrasting environments: Marine, terrestrial and human gut. Current Opinion in Microbiology, 70, 102229. 10.1016/j.mib.2022.102229

[2] Focardi, A., Ostrowski, M., Goossen, K., Brown, M. V., & Paulsen, I. (2020). Investigating the diversity of marine bacteriophage in contrasting water masses associated with the east australian current (EAC) system. Viruses, 12(3), 317. 10.3390/v12030317

[3] Naureen, Z., Dautaj, A., Anpilogov, K., Camilleri, G., Dhuli, K., Tanzi, B., Maltese, P. E., Cristofoli, F., De Antoni, L., Beccari, T., Dundar, M., & Bertelli, M. (2020). Bacteriophages presence in nature and their role in the natural selection of bacterial populations. Acta Bio Medica Atenei Parmensis, 91(13-S), e2020024. 10.23750/abm.v91i13-S.10819

[4] Kasman, L. M., & Porter, L. D. (2022, September 26). Bacteriophages. U.S. National Library of Medicine. https://www.ncbi.nlm.nih.gov/books/NBK493185/#:∼:text=Although%20bacteriophages%20cannot%20infect%20and,between%20pathogenic%20and%20nonpathogenic%20bacteria

[5] Schultz, J., Modolon, F., Rosado, A. S., Voolstra, C. R., Sweet, M., & Peixoto, R. S. (2022,,August 30). Methods and strategies to uncover coral-associated microbial dark matter. U.S. National Library of Medicine. https://pmc.ncbi.nlm.nih.gov/articles/PMC9426423/

[6] Ritchie, K. B. (2006, September 20). Regulation of microbial populations by coral surface mucus and mucus-associated bacteria. Inter-Research Science Center. https://www.int-res.com/articles/meps2006/322/m322p001.pdf

[7] Rickert, C. A., Lutz, T. M., Marczynski, M., & Lieleg, O. (2020, May 20). Several sterilization strategies maintain the functionality of mucin glycoproteins - rickert - 2020 - macromolecular bioscience - wiley online library. Wiley Online Library. 10.1002/mabi.202000090

[8] Rohwer, F., Barr, J. J., Auro, R., & Furlan, M. (2013, May 20). Bacteriophage adhering to mucus provide a non–host-derived immunity. PNAS. 10.1073/pnas.1305923110

[9] Clokie, M. R., Millard, A. D., Letarov, A. V., & Heaphy, S. (2011). Phages in nature. Bacteriophage, 1(1), 31–45. 10.4161/bact.1.1.14942

[10] Weitz, J. S., Stock, C. A., Wilhelm, S. W., Bourouiba, L., Coleman, M. L., Buchan, A., Follows, M. J., Fuhrman, J. A., Jover, L. F., Lennon, J. T., Middelboe, M., Sonderegger, D. L., Suttle, C. A., Taylor, B. P., Frede Thingstad, T., Wilson, W. H., & Eric Wommack, K. (2015). A multitrophic model to quantify the effects of marine viruses on microbial food webs and ecosystem processes. The ISME Journal, 9(6), 1352–1364. 10.1038/ismej.2014.220

[11] Koskella, B., & Brockhurst, M. A. (2014). Bacteria–phage coevolution as a driver of ecological and evolutionary processes in microbial communities. FEMS Microbiology Reviews, 38(5), 916–931. 10.1111/1574-6976.12072

[12] Garin-Fernandez, A., Pereira-Flores, E., Glöckner, F. O., & Wichels, A. (2018). The North Sea goes viral: Occurrence and distribution of North Sea bacteriophages. Marine Genomics, 41, 31–41. 10.1016/j.margen.2018.05.004

[13] Batinovic, S., Wassef, F., Knowler, S. A., Rice, D. T. F., Stanton, C. R., Rose, J., Tucci, J., Nittami, T., Vinh, A., Drummond, G. R., Sobey, C. G., Chan, H. T., Seviour, R. J., Petrovski, S., & Franks, A. E. (2019). Bacteriophages in Natural and Artificial Environments. Pathogens, 8(3). 10.3390/pathogens8030100

[14] McFall-Ngai, M. J. (2014). The Importance of Microbes in Animal Development: Lessons from the Squid-Vibrio Symbiosis. Annual Review of Microbiology, 68, 177–194. 10.1146/annurev-micro-091313-103654

[15] Wallace, B. A., Varona, N. S., Hesketh-Best, P. J., Stiffler, A. K., & Silveira, C. B. (2024). Globally distributed bacteriophage genomes reveal mechanisms of tripartite phage-bacteria-coral interactions. The ISME Journal, 18(1). 10.1093/ismejo/wrae132

[16] Thurber, R. V., Payet, J. P., Thurber, A. R., & Correa, A. M. S. (2017). Virus–host interactions and their roles in coral reef health and disease. Nature Reviews Microbiology, 15(4), 205–216. 10.1038/nrmicro.2016.176

[17] Roach T. N. F., Little, M., Arts, I., Huckeba, J., Haas, A. F., George, E. E., Quinn, R. A., Cobián-Güemes, A. G., Naliboff, D. S., Silveira, C. B., Mark, Linda Wegley Kelly, Dorrestein, P. C., & Rohwer, F. (2020). A multiomic analysis of in situ coral–turf algal interactions. Proceedings of the National Academy of Sciences of the United States of America, 117(24), 13588–13595. 10.1073/pnas.1915455117

[18] Rubio-Portillo, E., Robertson, S., & Antón, J. (2024). Coral mucus as a reservoir of bacteriophages targeting Vibrio pathogens. The ISME Journal, 18(1). 10.1093/ismejo/wrae017

[19] Cohen, Y., Joseph Pollock, F., Rosenberg, E., & Bourne, D. G. (2013). Phage therapy treatment of the coral pathogen Vibrio coralliilyticus. MicrobiologyOpen, 2(1), 64–74. 10.1002/mbo3.52

[20] Ling, H., Lou, X., Luo, Q., He, Z., Sun, M., & Sun, J. (2022). Recent advances in bacteriophage-based therapeutics: Insight into the post-antibiotic era. Acta Pharmaceutica Sinica B, 12(12). 10.1016/j.apsb.2022.05.007

[21] Küçükgülmez, A. (2018). Extraction of chitin from crayfish (Astacus leptodactylus) shell waste. Alinteri Zirai Bilimler Dergisi, 33(1), 99–104 https://experiments.springernature.com/articles/10.1007/978-1-0716-2601-6_11

[22] Liu, X., Zhang, M., Xie, Y., Zhang, L., & Ren, J. (2019). Biocompatibility, bioactivity, and therapeutic potential of nanocellulose-based hydrogels in biomedical applications. Frontiers in Bioengineering and Biotechnology, 7, 230. 10.3389/fbioe.2019.00230

[23] Draget, K. I., Skjåk-Bræk, G., & Smidsrød, O. (1994). Alginate based new materials. International Journal of Biological Macromolecules, 16(6), 305–309 10.1016/S0141-8130(97)00040-8

[24] Aiba, Y., Tan, X.-E., Veeranarayanan, S., Thitiananpakorn, K., Nguyen, H. M., & Wannigama, D. L. (2024). Phage therapy: A promising approach to combat multidrug-resistant bacteria. Antibiotics, 13(9), 870. 10.3390/antibiotics13090870

[25] Shapiro Karen, Krusor Colin, Mazzillo Fernanda F.M., Conrad Patricia A., Largier John L., Mazet Jonna A.K. and Silver Mary W. 2014 Aquatic polymers can drive pathogen transmission in coastal ecosystems Proc. R. Soc. B.28120141287. 10.1098/rspb.2014.1287

[26] Kauffman, K. M., Chang, W. K., Brown, J. M., Hussain, F. A., Yang, J., Polz, M. F., & Kelly, L. (2022). Resolving the structure of phage–bacteria interactions in the context of natural diversity. Nature Communications, 13(1). 10.1038/s41467-021-27583-z

[27] Echeverría-Vega, A., Morales-Vicencio, P., Saez-Saavedra, C., Gordillo-Fuenzalida, F., & Araya, R. (2019). A rapid and simple protocol for the isolation of bacteriophages from coastal organisms. MethodsX, 6, 2614–2619. 10.1016/j.mex.2019.11.003

